# Young Adults Use Whole-Body Feedback to Perceive Small Locomotor Disturbances

**DOI:** 10.1101/2022.09.08.507184

**Authors:** Daniel J. Liss, Hannah D. Carey, Jessica L. Allen

## Abstract

To prevent a fall when a disturbance to walking is encountered requires sensory information about the disturbance to be perceived, integrated, and then used to generate an appropriate corrective response. Prior research has shown that feedback of whole-body motion drives this corrective response. Here, we hypothesized that young adults also use whole-body motion to *perceive* locomotor disturbances. 15 subjects performed a locomotor discrimination task in which the supporting leg was slowed during stance every 8-12 steps to emulate subtle slips. The perception threshold of these disturbances was determined using a psychometrics approach and found to be 0.08 ± 0.03 m/s. Whole-body feedback was examined through center-of-mass (CoM) kinematics and whole-body angular momentum (WBAM). Perturbation-induced deviations of CoM and WBAM were calculated in response to the two perturbation levels nearest each subject’s perception threshold. Consistent with our hypothesis, we identified significantly higher perturbation induced deviations for perceived perturbations in sagittal-plane WBAM, anteroposterior CoM velocity, and mediolateral CoM position, velocity, and acceleration. Because whole body motion is not sensed directly but instead arises from the integration of various sensory feedback signals, we also explored local sensory feedback contributions to the perception of locomotor disturbances. Local sensory feedback was estimated through kinematic analogues of vision (head angle), vestibular (head angular velocity), proprioception (i.e., sagittal hip, knee, and ankle angles), and somatosensation (i.e., anterior-posterior & mediolateral center-of-pressure, COP). We identified significantly higher perturbation induced deviations for perceived perturbations in sagittal-plane ankle angle only. These results provide evidence for both whole-body feedback and ankle proprioception as important for the perception of subtle slip-like locomotor disturbances in young adults. Our interpretation is ankle proprioception is a dominant contributor to estimates of whole-body motion to perceive locomotor disturbances.

## 1. Introduction

Falls due to slips and trips are a leading cause of injuries (Cho et al., 2021; Heijnen & Rietdyk, 2016; Talbot et al., 2005). To prevent a fall when such a disturbance is encountered requires a rapid corrective response. Sensory information about the disturbance must be perceived, integrated, and used to generate an appropriate corrective response. We previously found that young adults can perceive subtle slip-like locomotor disturbances of less than 10% of their self-selected walking speed (Liss et al., 2022); however, we do not know what sources of sensory feedback contributed to this perceptual ability. This study therefore aims to understand what sensory feedback modalities contribute to the ability to perceive subtle slip-like locomotor disturbances in young adults. Understanding the role of sensory feedback in perceiving locomotor disturbances in young adults may have implications for populations with poor sensory feedback that are at a high risk of falls, e.g., older adults, persons with Parkinson’s disease, stroke survivors, etc.

Controlling whole-body motion is important to maintain balance while navigating the world. For example, keeping the center of mass (CoM) within the base of support is crucial for maintaining balance (Tesio & Rota, 2019). Whole-body angular momentum (WBAM) is also quickly controlled to prevent slips and trips (Pijnappels et al., 2004, 2005). Since controlling whole-body motion is crucial for balance, it is likely to also be sensed. Indeed, prior studies have found that feedback of CoM motion appears to drive muscle recruitment to maintain balance in response to both perturbations to walking and standing (Afschrift et al., 2021; Welch & Ting, 2009). However, it remains unknown if whole-body feedback is a key parameter underlying the perception of subtle slip-like locomotor disturbances.

Whole body motion is not sensed directly but instead arises from the integration of various sensory feedback signals, e.g., vision, vestibular, proprioception, somatosensation, etc. (Bent et al., 2004; Horak, 2006; Lephart et al., 1998; Peterka, 2002). Each of these sensory systems detect different types of information, which are then integrated to estimate where the body is in space (e.g., the proprioceptive system senses body and limb position whereas the vestibular system senses head orientation with respect to gravity (Peterka, 2002)). The relative importance of each system in estimating whole body motion can be modulated based on sensory context, e.g., increased reliance on vision when standing on different surfaces) (Lubetzky-Vilnai et al., 2015; Patel et al., 2008), threat to balance (e.g. standing at height) (Carpenter et al., 2001; Davis et al., 2009), phase of movement (e.g., modulation of somatosensory feedback in stance versus swing phase of walking) (Duysens et al., 1990; Perry et al., 2000), with loss or degradation of different sensory systems (e.g., increased reliance on vision in older adults with poor proprioception) (Franz & Thelen, 2015; Peterka, 2002), etc. The relative importance of these different sensory systems in perceiving subtle slip-like locomotor disturbances under normal conditions remains unknown.

This study aims to explore the role of whole-body and local sensory feedback on the perception of locomotor disturbances in healthy young adults. We hypothesized that young adults rely on whole-body sensory feedback to perceive locomotor disturbances. Based on this hypothesis, we predicted that perceived perturbations would have larger deviations in both CoM and WBAM than non-perceived perturbations. We also explored local sensory feedback contributions to the perception of locomotor disturbances, where local sensory feedback was estimated through kinematic analogues of vision (head angle), vestibular (head angular velocity), proprioception (i.e., sagittal hip, knee, and ankle angles), and somatosensation (i.e., anterior-posterior & mediolateral center-of-pressure, COP).

## 2. Methods

### 2.1 Subjects

Sixteen young adults (7 Females, 22.4 ± 2.80 years old) participated. All subjects were in good health with no self-reported history of recent lower extremity injuries, neurological deficits, vision, or balance problems. Subjects were novel to the experimental paradigm used in this study and provided written informed consent before participating according to the protocol approved by the Institutional Review Board of West Virginia University. Twelve out of sixteen subjects were the same subjects in a paper previously published in which we analyzed locomotor perception thresholds (Liss et al., 2022).

### 2.2 Experimental Protocol

Subjects wore a support harness while walking on a split-belt treadmill (Bertec, Columbus, OH) that collected ground reaction forces (GRFs) under each foot at 1000 Hz. Whole-body motion was recorded at 100 Hz using a ten-camera system with an extended lower-body plug-in gait marker set (Vicon Motion Systems, Oxford, UK) that included additional markers on the trunk and head. Subjects were allowed several minutes to adapt to treadmill walking before determining self-selected walking speed (SSWS). Briefly, SSWS was determined by asking if the subject was a “fast” or “slow” walker then moving to pace they would comfortably walk at for at least 10 min; a more detailed description can be found in our previous publication (Liss et al., 2022).

Subjects then performed the following locomotor discrimination task. Approximately every 8–12 steps, the supporting leg was slowed during stance (Fig. 1A). Three subjects also received accelerations on the supporting leg during stance, but only decelerations were used for analyses. The response to such locomotor perturbations induces similar kinematic responses as real-world slips and trips (Luukinen et al., 2000). The speed change was automatically triggered at heel-strike, defined as the point where the vertical GRF achieved 20% of the subject body weight. The perturbed belt was slowed by a magnitude (dV) of 0, 0.02, 0.05, 0.1, 0.15, 0.2, 0.3, and 0.4 m/s and returned to SSWS during the following swing phase (Fig. 1A). These eight dV disturbances were randomized and repeated five times per leg (80 total deceleration perturbations). Subjects were asked to respond “Yes” or “No” to whether they felt a disturbance to their balance after each perturbation, including catch trials of dV = 0. Subjects were instructed not to use safety rails for support or look at their feet and they wore noise-canceling headphones playing white noise to prevent auditory feedback of treadmill speed changes. The following measures were taken to increase consistency between subjects and prevent fatigue: all subjects received standardized instructions and the discrimination task was split into blocks of approximately 20 perturbations with at least 5 minutes of rest between blocks.

**Figure 1:**
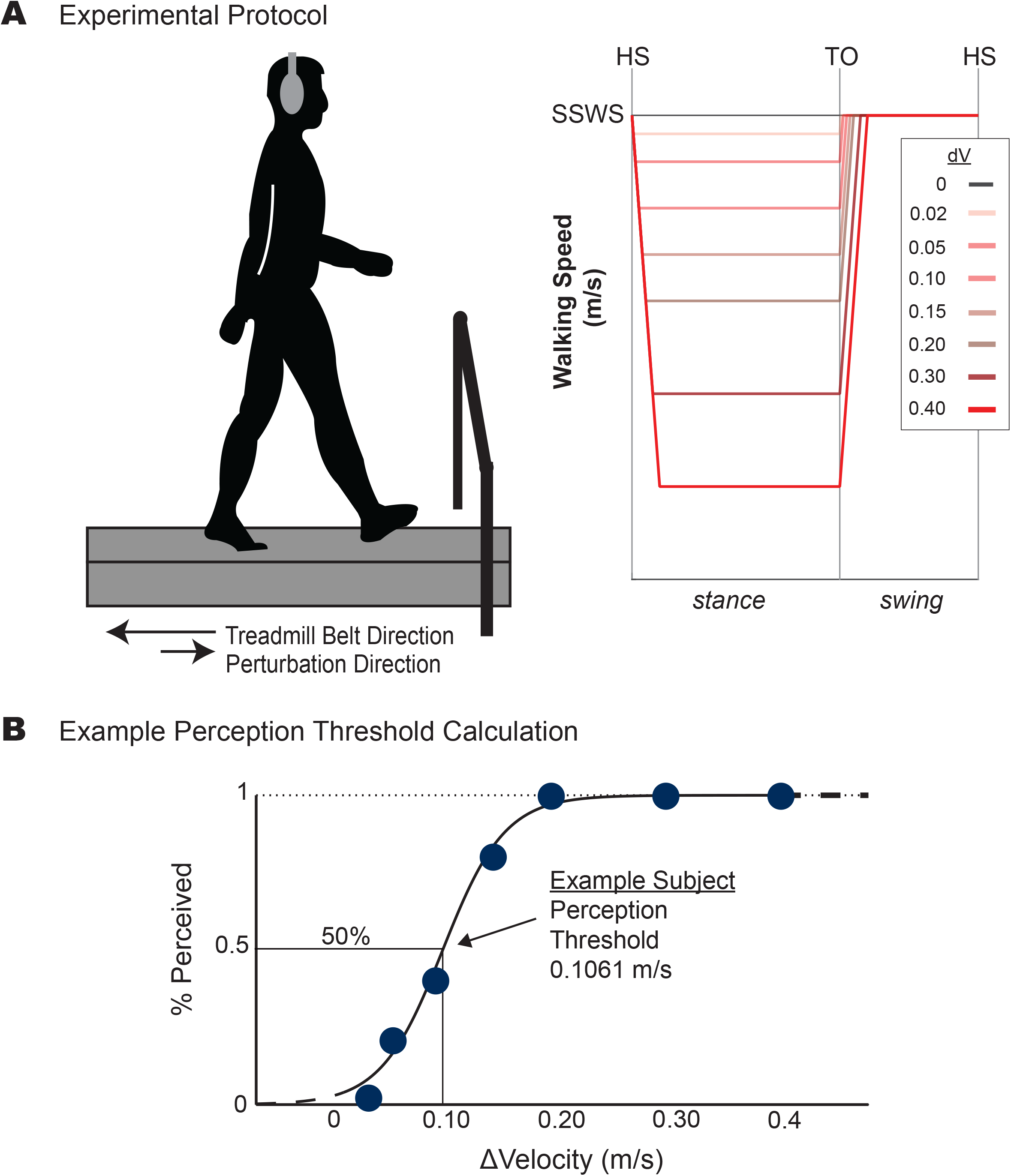
A) Subjects walked at their self-selected walking speed (SSWS) on a split-belt treadmill while receiving a subtle slip-like disturbance every 8-12 strides. Eight perturbation speeds (dV) were randomized and repeated 5 times on the dominant leg with perturbations on the nondominant leg random interspersed. All perturbations were triggered at heel-strike and the belt speed was returned to SSWS during the next swing phase. B) The perception threshold was determined as the 50% point (e.g., the just noticeable difference) of the psychometric fit to the percentage of perceived disturbances of each dV. In the example subject shown, the perception threshold was identified as 0.106 m/s.

### 2.3 Data Processing

#### 2.3.1 Disturbance Perception Threshold

The perception threshold was calculated for the dominant leg using the psignifit 4 MATLAB toolbox (Wichmann & Hill, 2001). Since we did not observe differences between legs in the perception threshold in our previous paper (Liss et al., 2022), we focused our analyses on the dominant leg. Leg dominance was labeled as the limb preferred when kicking a ball. The psignifit4 toolbox fits a logistical sigmoid curve to the proportion of perceived disturbances (out of 5 repetitions) across all perturbation magnitudes (Fig. 1B). The perception threshold was defined as 50% of the fit (i.e., the just noticeable difference). Because significant guessing (i.e., positive “Yes” responses at the 0 m/s disturbance) prevents accurate identification of this threshold, we removed any subject with a positive response in more than one-third of trials with dV=0 (i.e., the catch trial). Based on this cutoff, one subject was removed from further analyses.

#### 2.3.2 Whole-body and local sensory feedback calculations

All subjects had a 3D OpenSim model (gait2392) scaled to their anthropometrics using the ‘Scale Tool’ (Delp et al., 2007) based on markers placed on or near bony landmarks recorded during a 4-5 second static calibration trial (Table 1).

**Table 1:**
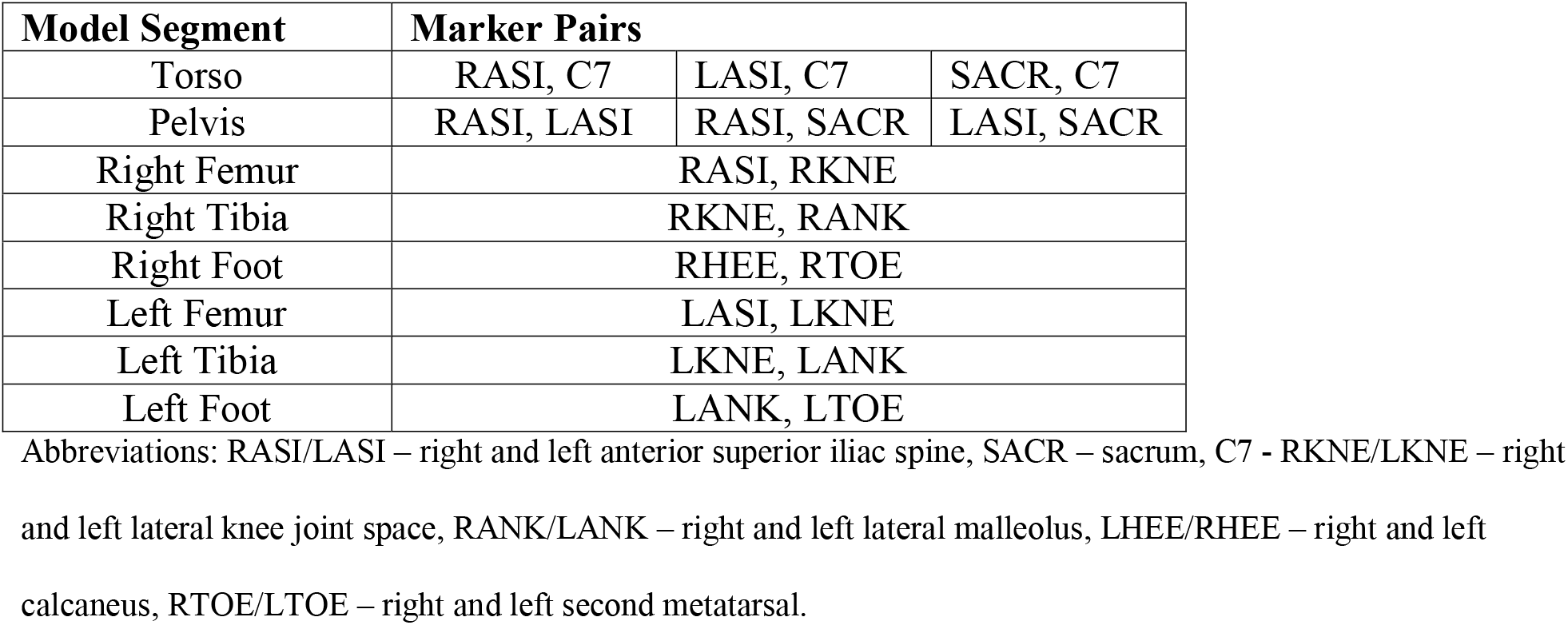
Marker pairs used to scale an OpenSim gait2392 model for each subject.

The scaled model was combined with marker data for each walking trial to calculate joint angles using the ‘Inverse Kinematics Tool’ and body segment CoM kinematics using the ‘Body Analysis Tool’. All data was then split up by gait cycle defined from heel strike to heel strike and normalized to 101 data points (representing 0 – 100% of the gait cycle at 1% increments).

Whole-body feedback variables included CoM and WBAM. CoM position was calculated using the weighted sum of all the model’s individual segment masses (Winter, 2009). CoM velocity and acceleration were calculated as the first and second derivatives of CoM position, respectively (Fig. 2A-C). For position, we focused our analysis on CoM position *deviation* by subtracting the position at heel-strike for each gait cycle to remove bias from subjects wandering on the treadmill.

**Figure 2:**
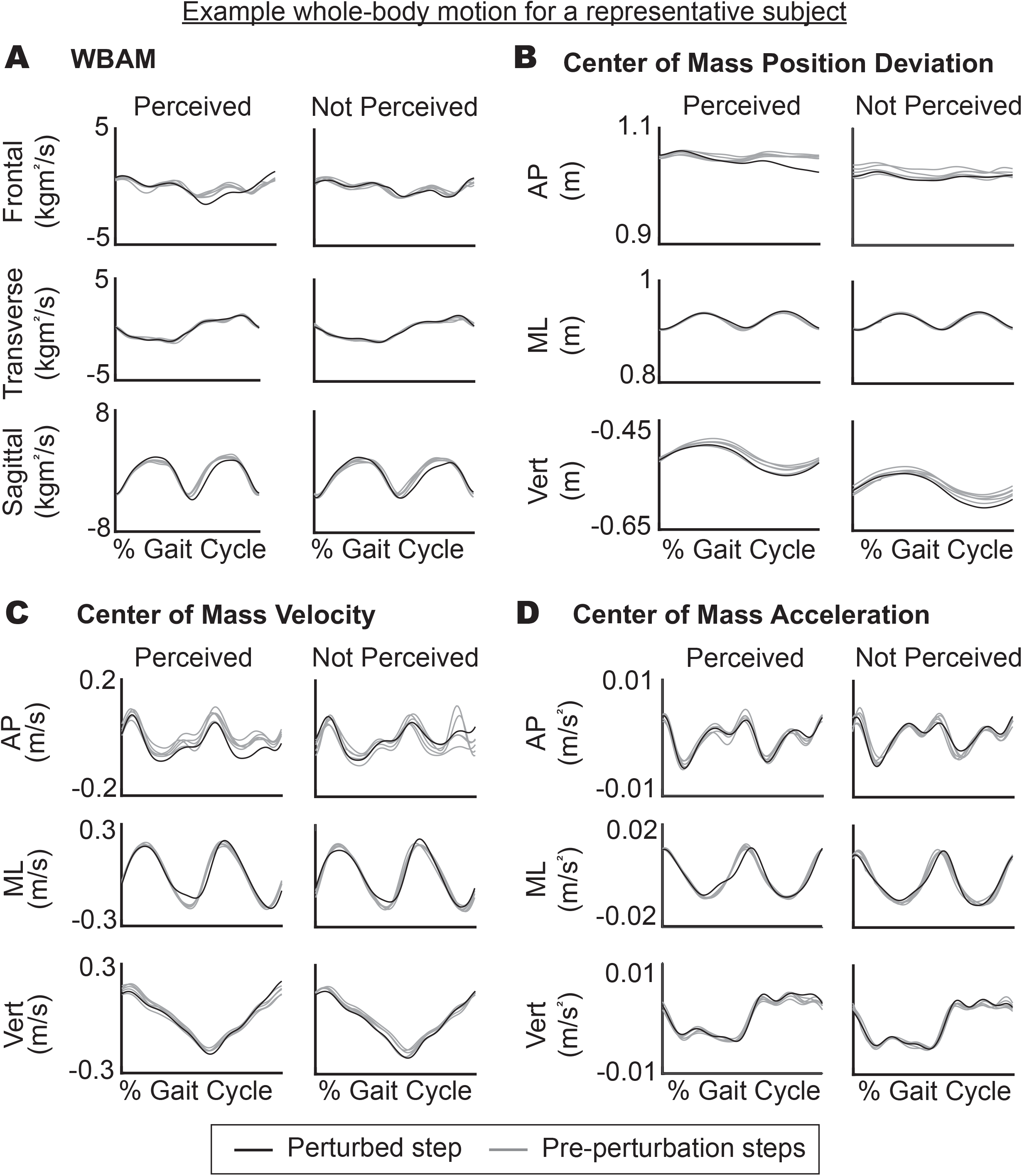
Representative subject demonstrating differences between perceived and not perceived perturbations for A) Whole-body angular momentum (WBAM), B) Center-of-mass (CoM) position, C) CoM velocity, and D) CoM acceleration. Representative subject is the same as the perception curve in Figure 1C, with curves taken from 0.10 m/s perturbations just below their perception threshold. (Gray - 5 gait cycles before the perturbation; Black - perturbed gait cycle).

Whole-body angular momentum in each plane of motion was calculated (Fig. 2D) using the kinematic equation from Herr & Popovic, 2008:

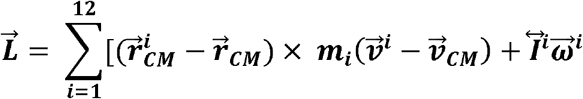

where ***i*** is model segment (i.e., torso, femur, etc.), 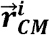 is CoM position for each model segment, 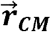 is the overall body CoM position, **m_i_** is the mass of each model segment, 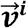 is CoM velocity of each model segment, 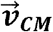 is the velocity of the overall body CoM, 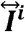 is the inertia tensor for each model segment, and 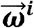 is the angular velocity of each model segment.

For local sensory feedback analogues, head angle (visual analogue), head angular velocity (vestibular analogue), sagittal hip, knee, and ankle angles (proprioceptive analogue), and anterior-posterior center-of-pressure (COP, somatosensation analogue) were used for analysis. All analyses focused on the dominant leg. Head calculations (i.e., sagittal plane head angle and head angular velocity) were calculated by forming a plane from the 4 head markers worn during the experiment (Fig. 3A&B). Sagittal hip, knee, and ankle angles were taken from the inverse kinematics results from OpenSim (Fig. 3C). COP was collected from the Bertec instrumented treadmill force plates and analyzed over stance phase only (Fig. 3D).

**Figure 3:**
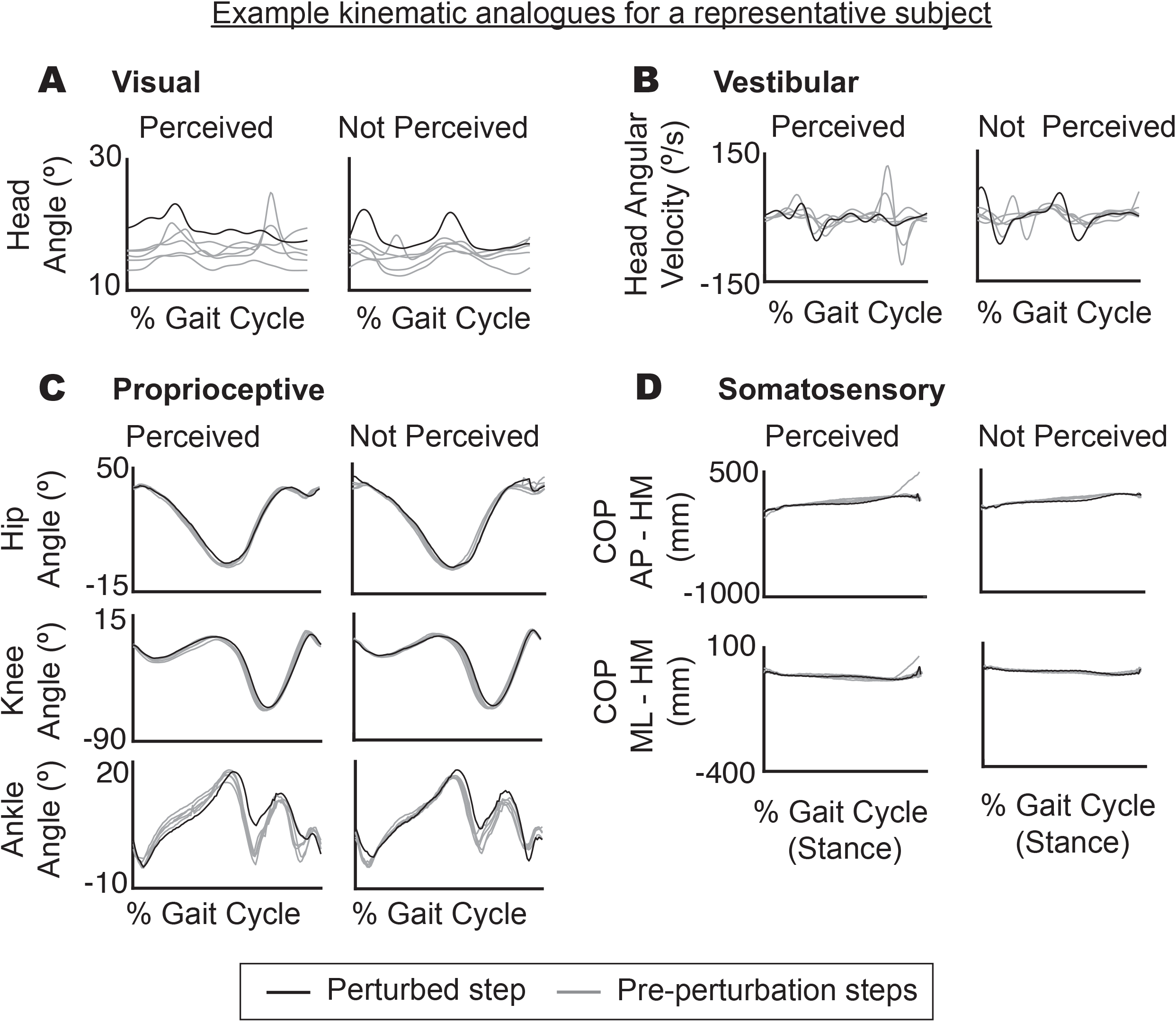
Representative subject demonstrating differences in kinematic analogues of local sensory feedback in perceived versus not perceived perturbations. A) Visual feedback was estimated by sagittal head angle. B) Vestibular feedback was estimated by sagittal head angular velocity. C) Proprioceptive feedback was estimated by sagittal hip, knee, and angle angles. D) Somatosensory feedback was estimated through center of pressure (AP & ML) minus the heel marker of the dominant foot in the respective direction. Representative subject is the same as the perception curve in Figure 1C, with curves taken from 0.10 m/s perturbations just below their perception threshold. (Gray - 5 gait cycles before the perturbation; Black - perturbed gait cycle).

#### 2.3.3 Perturbation-Induced Deviation Calculation

Perturbation-induced deviations of each whole body and local sensory feedback variable were calculated in response to the two perturbation levels nearest each subject’s perception threshold. For example, for a subject whose threshold was 0.12 m/s we analyzed the trials for dV = 0.10 m/s and dV = 0.15 m/s. This resulted in 10 trials analyzed per subject. For each variable, the deviation between the perturbed gait cycle and the five gait cycles immediately preceding the perturbation was calculated as

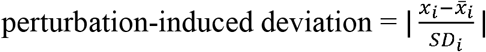

where i is % gait cycle, x is the perturbed gait cycle, and 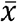 and SD are the average and standard deviation of the pre-perturbation gait cycles (see Fig. 4 for an example). All deviations were then averaged over the gait cycle to calculate a single value for each perturbation.

**Figure 4:**
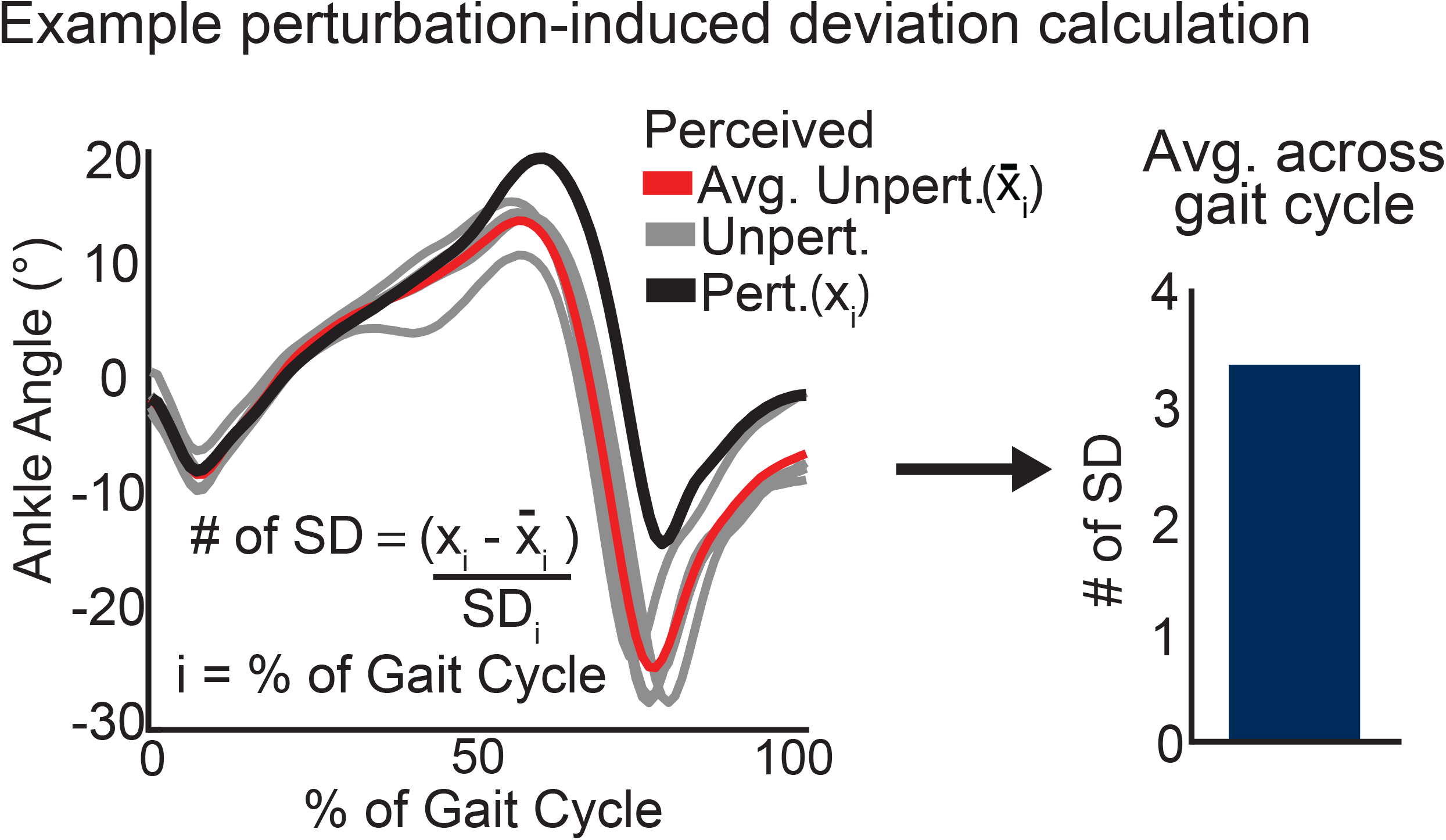
Perturbation-induced deviation was calculated for each whole-body and local feedback parameter (from Fig. 2 and 3). This was calculated at each percentage of the gait cycle, *i*, by taking the perturbed cycle, *x*, and subtracting the average of the five pre-perturbation gait cycles, 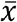, and then dividing by the standard deviation of the pre perturbation cycles, *SD* (Left). This value was then averaged over the gait cycle to get a single value for each perturbation (Right). (Black − perturbed gait cycle; Gray – 5 pre-perturbation gait cycles; Red – average of the pre-perturbation gait cycles).

#### 2.3.4 Statistical Analysis

All statistical analyses were performed in MATLAB (The MathWorks, Inc., Natick, Massachusetts, United States). To test our prediction that perceived perturbations would have larger deviations in CoM kinematics than non-perceived perturbations, we used separate Wilcoxon rank sum tests for each CoM kinematic variable (position, velocity and acceleration in all 3 directions = 9 variables) with Bonferroni corrections to correct for multiple comparisons (α=0.05/9 CoM variables = 0.0055). Similarly, we used a separate set of Wilcoxon rank sums tests to test our prediction that perceived perturbations would have larger deviations in WBAM than non-perceived perturbations with Bonferroni corrections for multiple comparisons (α=0.05/3 planes = 0.017). Finally, we used another set of Wilcoxon rank sums test to explore which local sensory feedback variables differed between perceived and not perceived perturbations with Bonferroni correction for multiple comparisons (α=0.05/7 kinematic variables = 0.007). 20/150 perturbations couldn’t be analyzed for head angle and head angular velocity due to missing head markers.

As an additional analysis to explore the strength of the relationship between sensory feedback and perturbation perception, individual logistical mixed-effect models were run for each of the significant parameters from the whole-body and local sensory feedback comparisons as described above. Each model treated the significant variable (*X*) as the fixed-effect and subject (*g*) as a random effect: *perceived* ~ *X* + (*X* | *g*). The MATLAB function ‘fitglm’ with binomial distribution and logit link function was used.

## 3. Results

Subjects walked at a SSWS of 1.14 ± 0.11 m/s and only one subject was left leg dominant. Subjects perceived locomotor disturbances of 0.08 ± 0.03 m/s on their dominant leg and there was no significant correlation between SSWS and perception threshold (p = 1.00).

WBAM and CoM kinematics were analyzed to examine contributions to locomotor perception from whole-body feedback. For angular momentum, only sagittal WBAM was significantly higher for perceived versus not perceived perturbations (frontal, p = 0.047; transverse, p = 0.127; sagittal, p = 0.005; with α = 0.05/3 = 0.017, Fig. 5A). For CoM kinematics, there were significantly higher perturbation-induced deviations in perceived perturbations for position, velocity, and acceleration. However, CoM position deviation significantly differed only in the mediolateral direction (AP, p = 0.02; ML, p = 0.003, Vertical, p = 0.035; with α = 0.05/9 = 0.0055, Fig. 5B). Similarly, CoM acceleration was only significantly different in the mediolateral direction (AP, p = 0.30; ML, p = 0.005; Vertical, p = 0.144, Fig. 5B) whereas CoM velocity significantly differed in both the AP (p = 0.001) and ML (p = 0.001) directions but not the vertical direction (p = 0.03, Fig. 5B).

**Figure 5:**
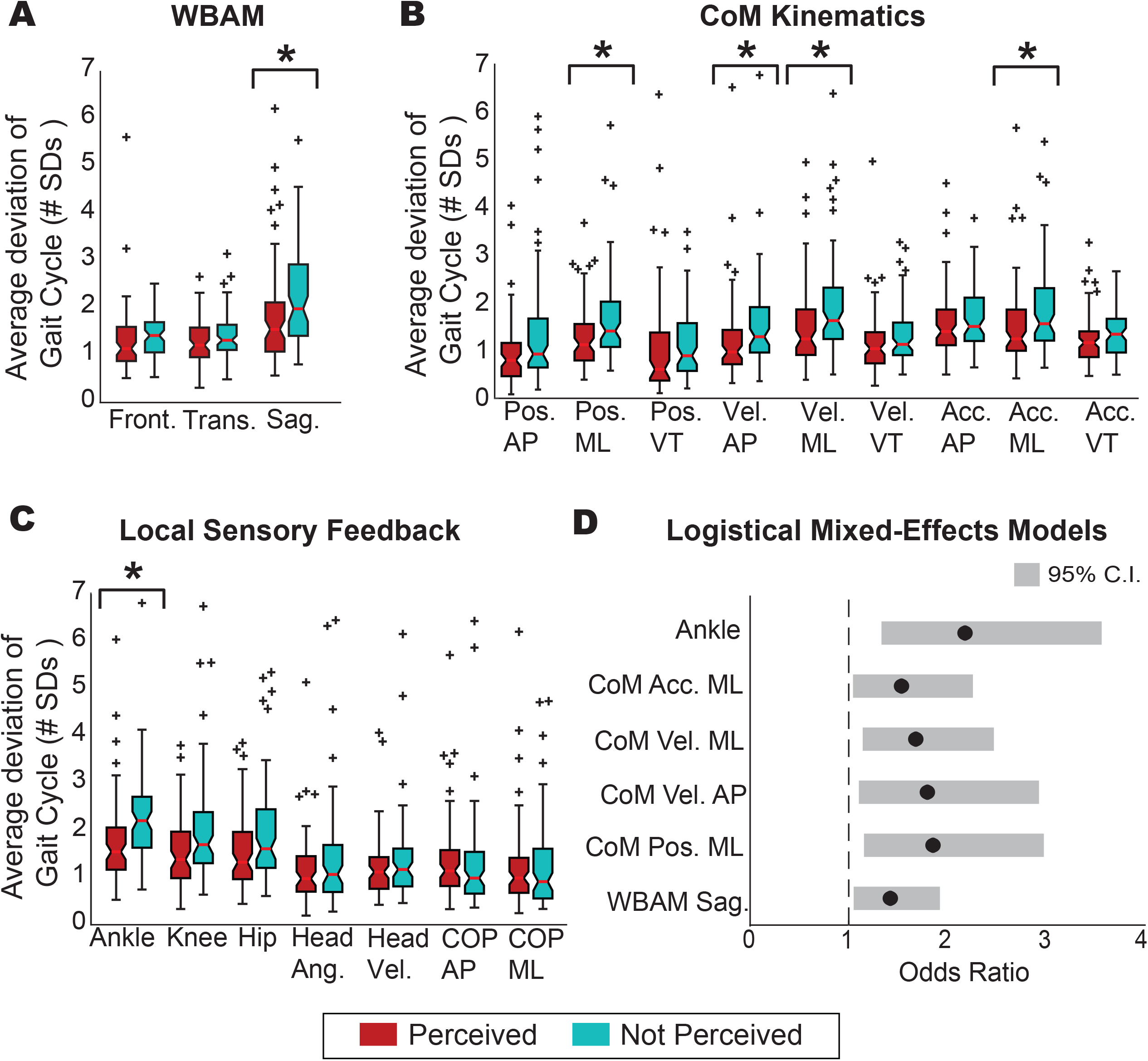
Differences in perceived versus not perceived perturbations for A: whole-body angular momentum (WBAM), B: center-of-mass (CoM) kinematics, and C) local sensory feedback analogues. A) Only sagittal plane WBAM was significantly higher in perceived versus not perceived perturbations. B) ML CoM position deviation, AP & ML CoM Velocity, and ML CoM acceleration were all significantly larger in perceived versus not perceived perturbations. C) Ankle proprioception was the only local sensory feedback variable that was significantly larger in perceived versus not perceived trials. D) The significant variables identified in A,B, and C were run through individual logistical mixed-effects models. The lower bound of the 95% confidence interval of the odds ratio for each variable did not cross one, suggesting a significant predictive effect of perturbation induced deviations and the perception of a disturbance. (*signifies a p value < Bonferroni corrected α value.)

Local sensory feedback analogues were analyzed to examine contributions from separate sensory systems. Visual (head angle, p = 0.53, Fig. 5C), vestibular (head angular velocity, p = 0.26, Fig. 5C), and somatosensory (COP-AP, p = 0.30 and COP-ML, p = 0.60, Fig. 5C) sensory feedback analogues did not significantly differ between perceived versus not perceived perturbations. For the proprioceptive analogues, only ankle angle deviation (p < 0.001 with α = 0.05/7 = 0.007, Fig. 5C) was significantly higher for perceived perturbations (knee angle deviation, p =0.008; hip angle deviation, p = 0.017, Fig. 5C).

Individual logistical mixed-effect models were run on the significant parameters from both whole-body and local sensory feedback (i.e., WBAM sagittal, CoM ML position, CoM AP & ML velocity, CoM ML acceleration, and ankle angle). All models were significant (i.e., the 95% confidence interval of their odds ratio did not cross 1, Fig. 5D) with no random effect of subject. WBAM sagittal had an odds ratio of 1.4, CoM ML position deviation had an odds ratio of 1.9, CoM AP velocity had an odds ratio of 1.8, CoM ML velocity had an odds ratio 1.7, CoM ML acceleration had an odds ratio of 1.5, and ankle angle had an odds ratio of 2.2.

## 4. Discussion

This study examined the role whole-body and local sensory feedback play in perceiving subtle slip-like locomotor disturbances in young adults. Based on our hypothesis that young adults would rely on whole-body feedback to perceive disturbances, we predicted that perceived perturbations would have larger deviations in CoM and WBAM than non-perceived perturbations. This prediction was supported, providing evidence for whole-body feedback as important for the perception of subtle slip-like locomotor disturbances. We also explored local sensory feedback contributions and found that only perturbation-induced deviations in ankle angle differed between perceived versus not perceived perturbations. These results support the important role ankle proprioception may play in perceiving subtle slip and trip-like locomotor disturbances.

### 4.1 Whole-body feedback is important for perceiving locomotor disturbances

Our results support the role of whole-body feedback for maintaining balance in challenging environments. In particular, we found that sensing whole-body motion may contribute to the perception of slip-like locomotor disturbances. Moreover, these results point to the importance of both translational (CoM kinematics) and rotational (WBAM) whole-body sensing for locomotor disturbance perception.

Consistent with our predictions, we found that perturbation induced CoM kinematics were significantly larger in perceived versus not perceived perturbations. Interestingly, we found differences between perceived and not perceived perturbations in both the AP and ML directions despite the belt speed changes being a primarily AP disturbance. The change in AP CoM kinematics is consistent with prior studies demonstrating changes in the AP direction in response to AP perturbations (Afschrift et al., 2021; Vlutters et al., 2016; Welch & Ting, 2009). That we also found differences in ML CoM kinematics is consistent with humans being laterally unstable during walking (Kuo, 1999; McAndrew et al., 2011) and increasing their step width in response to AP perturbations to maintain stability (Dean et al., 2007; Golyski et al., 2022; Roeles et al., 2018). We expand on these prior findings by demonstrating that within perturbations of *similar magnitude*, differences in perturbation induced CoM kinematics may explain whether a perturbation was perceived or not. Taken together with prior studies that have identified feedback of CoM kinematics as a driver of muscle recruitment (Afschrift et al., 2021; Welch & Ting, 2009, 2014), there is strong evidence for the importance of whole-body CoM kinematics for the perception of locomotor disturbances.

We also provide evidence that in addition to linear CoM motion we may monitor rotational motion about the CoM. Consistent with our predictions, we found larger perturbation induced deviations in WBAM about the CoM in perceived versus not perceived disturbances, but only in the sagittal plane (Fig. 5A). Classically, WBAM about the CoM has been used to describe the state of dynamic balance (e.g., impaired versus not impaired populations (Honda et al., 2019; Nott et al., 2014), across different tasks (Silverman et al., 2012, 2014), etc.). Prior studies have identified that perturbations emulating slips and trips lead to higher WBAM compared to normal walking in healthy young adults (Liu et al., 2018; Martelli et al., 2013; Pijnappels et al., 2005; Potocanac et al., 2014). In agreement with these prior studies, post-hoc analyses in the current study revealed that WBAM was higher in all perturbations (both perceived and not-perceived) compared to non-perturbed walking (p<0.0001 for all three planes). While prior studies examining dynamic balance have focused on relatively large disturbances (e.g., 50% of walking speed (Liu et al., 2018) 100% of walking speed (Golyski et al., 2022),, 150% of walking speed (Martelli et al., 2013)), our results show that measurable differences in WBAM also occur for small perturbations (i.e., < 10% of walking speed). Moreover, our primary analyses identified significantly larger perturbation induced deviations in sagittal plane WBAM in perceived versus not perceived disturbances despite the perturbations being of similar magnitude. Such a result suggest that we may be sensitive to angular motion about the CoM when perceiving if a disturbance has occurred or not. Although we did not identify increases in frontal and transverse plane WBAM associated with perceived trials in the current study, we expect that perturbations with more mediolateral components would reveal that we monitor angular motion about all planes of movement.

### 4.2 Ankle proprioception is important for perceiving slip-like locomotor disturbances

Consistent with prior literature demonstrating the importance of ankle proprioception for locomotor control, our results provide evidence that ankle proprioception is also important for the perception of locomotor disturbances. Prior studies have primarily focused on proprioceptive acuity of either passive or active motion during sitting or standing and identified that worse proprioceptive acuity at the ankle is associated with worse self-reported and clinical measures of balance and walking in balance-impaired individuals (e.g., older adults, (Deshpande et al., 2016; Westlake et al., 2007)). Here, we expand upon those results to examine ankle proprioception during walking and in younger adults without balance impairments. We found that our kinematic analogue of ankle proprioception was associated with locomotor disturbance perception such that perceived disturbances had higher perturbation induced deviation in ankle angle (Fig. 5C). Moreover, for every 1 standard deviation increase in ankle angle during the perturbed gait cycle compared to the preceding steps, subjects were over two times more likely to perceive that a perturbation had occurred (Fig. 5D). These results support the importance of ankle proprioception and may help explain why poor distal proprioception has been associated with worse balance control (Deshpande et al., 2016; Westlake et al., 2007). As proprioception deteriorates there is often an increased reliance on visual and/or vestibular feedback. Given that the subtle slips perceived by young adults did not elicit significant differences in our visual and vestibular kinematic analogues, increased reliance on these sensory feedback modalities would likely require larger perturbations in order to be perceived.

Whether ankle proprioception directly led to disturbance perception or was instead the dominant contributor to estimates of whole-body motion that drove disturbance perception remains an open question. On the one hand, our kinematic analogue of ankle proprioception had the strongest relationship with disturbance perception out of all parameters (i.e., highest odds ratio, Fig. 5D). On the other hand, prior literature provides strong evidence that whole-body feedback and not local feedback drives the motor response to perturbations (Afschrift et al., 2021; Welch & Ting, 2014). For example, similar ankle angle changes are evoked in both rotational and translational surface perturbations to standing yet they produce opposite changes in CoM motion that require different motor responses (Nashner, 1976). Several more recent studies have demonstrated that muscle activity is better explained by feedback of whole-body motion than local joint angle changes (Afschrift et al., 2021; Welch & Ting, 2014). Moreover, even when ankle motion is restricted by wearing a boot there is still a corrective motor response in the ankle muscles in response to perturbations to walking (Vlutters et al., 2018). Taken together, these results suggest that ankle proprioception may be the dominant contributor to whole body estimates. However, just because whole-body feedback drives the motor response does not rule out that disturbance perception is driven by local sensory feedback. Future studies are needed to unravel the role of whole-body and local sensory feedback in both perception and action.

Although our kinematic analogue of ankle angle proprioception emerged as important for the subtle slip-like locomotor disturbances of the current study, it is likely that other joints, planes of movement, and sensory modalities may play an important role in different types of perturbations (e.g., waist-pulls, mediolateral slips and trips, etc.). However, it is important to note that we may have underestimated the role other sensory modalities play in perceiving subtle slip-like locomotor disturbances since we could not directly measure sensory feedback. Although our novel kinematics analogues allowed us to overcome this limitation to estimate local sensory feedback, future research should further evaluate the utility of these methods (e.g., against proprioceptive acuity tests, eye-tracking, galvanic stimulation, etc.).

## 5. Conclusion

We found that young adults may use whole-body feedback to perceive subtle slip-like locomotor disturbances. Our results also point to the importance of ankle proprioception in perceiving locomotor disturbances. This study serves as an important step to understand the role of sensory feedback in perceiving locomotor disturbances. Future work should investigate how this perceptual ability changes in individuals that have increased fall risk (e.g., due to musculoskeletal diseases, neurodegeneration, aging, etc.).

## References

Afschrift, M., De Groote, F., & Jonkers, I. (2021). Similar sensorimotor transformations control balance during standing and walking. PLOS Computational Biology, 17(6), e1008369. https://doi.org/10.1371/journal.pcbi.1008369

Bent, L. R., Inglis, J. T., & McFadyen, B. J. (2004). When is Vestibular Information Important During Walking? Journal of Neurophysiology, 92(3), 1269–1275. https://doi.org/10.1152/jn.01260.2003

Carpenter, M., Frank, J., Silcher, C., & Peysar, G. (2001). The influence of postural threat on the control of upright stance. Experimental Brain Research, 138(2), 210–218. https://doi.org/10.1007/s002210100681

Cho, H., Heijnen, M. J. H., Craig, B. A., & Rietdyk, S. (2021). Falls in young adults: The effect of sex, physical activity, and prescription medications. PLOS ONE, 16(4),e0250360. https://doi.org/10.1371/journal.pone.0250360

Davis, J. R., Campbell, A. D., Adkin, A. L., & Carpenter, M. G. (2009). The relationship between fear of falling and human postural control. Gait & Posture, 29(2), 275–279. https://doi.org/10.1016/j.gaitpost.2008.09.006

Dean, J. C., Alexander, N. B., & Kuo, A. D. (2007). The Effect of Lateral Stabilization on Walking in Young and Old Adults. IEEE Transactions on Biomedical Engineering, 54(11), 1919–1926. https://doi.org/10.1109/TBME.2007.901031

Delp, S. L., Anderson, F. C., Arnold, A. S., Loan, P., Habib, A., John, C. T., Guendelman, E., & Thelen, D. G. (2007). OpenSim: Open-Source Software to Create and Analyze Dynamic Simulations of Movement. IEEE Transactions on Biomedical Engineering, 54(11), 1940–1950. https://doi.org/10.1109/TBME.2007.901024

Deshpande, N., Simonsick, E., Metter, E. J., Ko, S., Ferrucci, L., & Studenski, S. (2016). Ankle proprioceptive acuity is associated with objective as well as self-report measures of balance, mobility, and physical function. AGE, 38(3), 53. https://doi.org/10.1007/s11357-016-9918-x

Duysens, J., Tax, A. A. M., Nawijn, S., Berger, W., Prokop, T., & Altenmüller, E. (1990). Gating of sensation and evoked potentials following foot stimulation during human gait. Experimental Brain Research, 105(3). https://doi.org/10.1007/BF00233042

Franz, J. R., & Thelen, D. G. (2015). Depth-dependent variations in Achilles tendon deformations with age are associated with reduced plantarflexor performance during walking. Journal of Applied Physiology, 119(3), 242–249. https://doi.org/10.1152/japplphysiol.00114.2015

Golyski, P. R., Vazquez, E., Leestma, J. K., & Sawicki, G. S. (2022). Onset timing of treadmill belt perturbations influences stability during walking. Journal of Biomechanics, 130, 110800. https://doi.org/10.1016/j.jbiomech.2021.110800

Heijnen, M. J. H., & Rietdyk, S. (2016). Falls in young adults: Perceived causes and environmental factors assessed with a daily online survey. Human Movement Science, 46, 86–95. https://doi.org/10.1016/j.humov.2015.12.007

Herr, H., & Popovic, M. (2008). Angular momentum in human walking. Journal of Experimental Biology, 211(4), 467–481. https://doi.org/10.1242/jeb.008573

Honda, K., Sekiguchi, Y., Muraki, T., & Izumi, S.-I. (2019). The differences in sagittal plane whole-body angular momentum during gait between patients with hemiparesis and healthy people. Journal of Biomechanics, 86, 204–209. https://doi.org/10.1016/j.jbiomech.2019.02.012

Horak, F. B. (2006). Postural orientation and equilibrium: What do we need to know about neural control of balance to prevent falls? Age and Ageing, 35(suppl_2), ii7–ii11. https://doi.org/10.1093/ageing/afl077

Kuo, A. D. (1999). Stabilization of Lateral Motion in Passive Dynamic Walking. The International Journal of Robotics Research, 18(9), 917–930. https://doi.org/10.1177/02783649922066655

Lephart, S. M., Pincivero, D. M., & Rozzi, S. L. (1998). Proprioception of the Ankle and Knee: Sports Medicine, 25(3), 149–155. https://doi.org/10.2165/00007256-199825030-00002

Liss, D. J., Carey, H. D., Yakovenko, S., & Allen, J. L. (2022). Young adults perceive small disturbances to their walking balance even when distracted. Gait & Posture, 91, 198–204. https://doi.org/10.1016/j.gaitpost.2021.10.019

Liu, C., Macedo, L. D., & Finley, J. M. (2018). Conservation of Reactive Stabilization Strategies in the Presence of Step Length Asymmetries During Walking. Frontiers in Human Neuroscience, 12, 251. https://doi.org/10.3389/fnhum.2018.00251

Lubetzky-Vilnai, A., McCoy, S. W., Price, R., & Ciol, M. A. (2015). Young Adults Largely Depend on Vision for Postural Control When Standing on a BOSU Ball but Not on Foam. Journal of Strength and Conditioning Research, 29(10), 2907–2918. https://doi.org/10.1519/JSC.0000000000000935

Luukinen, H., Herala, M., Koski, K., Honkanen, R., Laippala, P., & Kivelä, S.-L. (2000). Fracture Risk Associated with a Fall According to Type of Fall Among the Elderly. Osteoporosis International, 11(7), 631–634. https://doi.org/10.1007/s001980070086

Martelli, D., Monaco, V., Luciani, L. B., & Micera, S. (2013). Angular Momentum During Unexpected Multidirectional Perturbations Delivered While Walking. IEEE Transactions on Biomedical Engineering, 60(7), 1785–1795. https://doi.org/10.1109/TBME.2013.2241434

McAndrew, P. M., Wilken, J. M., & Dingwell, J. B. (2011). Dynamic stability of human walking in visually and mechanically destabilizing environments. Journal of Biomechanics, 44(4), 644–649. https://doi.org/10.1016/j.jbiomech.2010.11.007

Nashner, L. M. (1976). Adapting reflexes controlling the human posture. Experimental Brain Research, 26(1). https://doi.org/10.1007/BF00235249

Nott, C. R., Neptune, R. R., & Kautz, S. A. (2014). Relationships between frontal-plane angular momentum and clinical balance measures during post-stroke hemiparetic walking. Gait & Posture, 39(1), 129–134. https://doi.org/10.1016/j.gaitpost.2013.06.008

Patel, M., Fransson, P. A., Lush, D., & Gomez, S. (2008). The effect of foam surface properties on postural stability assessment while standing. Gait & Posture, 28(4), 649–656. https://doi.org/10.1016/j.gaitpost.2008.04.018

Perry, S. D., McIlroy, W. E., & Maki, B. E. (2000). The role of plantar cutaneous mechanoreceptors in the control of compensatory stepping reactions evoked by unpredictable, multi-directional perturbationq. Brain Research, 6.

Peterka, R. J. (2002). Sensorimotor Integration in Human Postural Control. Journal of Neurophysiology, 88(3), 1097–1118. https://doi.org/10.1152/jn.2002.88.3.1097

Pijnappels, M., Bobbert, M. F., & Dieën, J. H. van. (2005). Push-off reactions in recovery after tripping discriminate young subjects, older non-fallers and older fallers. Gait & Posture, 21(4), 388–394. https://doi.org/10.1016/j.gaitpost.2004.04.009

Pijnappels, M., Bobbert, M. F., & van Diee, J. H. (2004). Contribution of the support limb in control of angular momentum after tripping. Journal of Biomechanics, 8.

Potocanac, Z., de Bruin, J., van der Veen, S., Verschueren, S., van Dieën, J., Duysens, J., & Pijnappels, M. (2014). Fast online corrections of tripping responses. Experimental Brain Research, 232(11), 3579–3590. https://doi.org/10.1007/s00221-014-4038-2

Roeles, S., Rowe, P. J., Bruijn, S. M., Childs, C. R., Tarfali, G. D., Steenbrink, F., & Pijnappels, M. (2018). Gait stability in response to platform, belt, and sensory perturbations in young and older adults. Medical & Biological Engineering & Computing, 56(12), 2325–2335. https://doi.org/10.1007/s11517-018-1855-7

Silverman, A. K., Neptune, R. R., Sinitski, E. H., & Wilken, J. M. (2014). Whole-body angular momentum during stair ascent and descent. Gait & Posture, 39(4), 1109–1114. https://doi.org/10.1016/j.gaitpost.2014.01.025

Silverman, A. K., Wilken, J. M., Sinitski, E. H., & Neptune, R. R. (2012). Whole-body angular momentum in incline and decline walking. Journal of Biomechanics, 45(6), 965–971. https://doi.org/10.1016/j.jbiomech.2012.01.012

Talbot, L. A., Musiol, R. J., Witham, E. K., & Metter, E. J. (2005). Falls in young, middle-aged and older community dwelling adults: Perceived cause, environmental factors and injury. BMC Public Health, 5(1), 86. https://doi.org/10.1186/1471-2458-5-86

Tesio, L., & Rota, V. (2019). The Motion of Body Center of Mass During Walking: A Review Oriented to Clinical Applications. Frontiers in Neurology, 10, 999. https://doi.org/10.3389/fneur.2019.00999

Vlutters, M., Van Asseldonk, E. H. F., & Van der Kooij, H. (2016). Center of mass velocity based predictions in balance recovery following pelvis perturbations during human walking. Journal of Experimental Biology, jeb.129338. https://doi.org/10.1242/jeb.129338

Vlutters, M., van Asseldonk, E. H. F., & van der Kooij, H. (2018). Reduced center of pressure modulation elicits foot placement adjustments, but no additional trunk motion during anteroposterior-perturbed walking. Journal of Biomechanics, 68, 93–98. https://doi.org/10.1016/j.jbiomech.2017.12.021

Welch, T. D. J., & Ting, L. H. (2009). A Feedback Model Explains the Differential Scaling of Human Postural Responses to Perturbation Acceleration and Velocity. Journal of Neurophysiology, 101(6), 3294–3309. https://doi.org/10.1152/jn.90775.2008

Welch, T. D. J., & Ting, L. H. (2014). Mechanisms of Motor Adaptation in Reactive Balance Control. PLoS ONE, 9(5), e96440. https://doi.org/10.1371/journal.pone.0096440

Westlake, K. P., Wu, Y., & Culham, E. G. (2007). Velocity discrimination: Reliability and construct validity in older adults. Human Movement Science, 26(3), 443–456. https://doi.org/10.1016/j.humov.2006.12.002

Wichmann, F. A., & Hill, N. J. (2001). The psychometric function: I. Fitting, sampling, and goodness of fit. Perception & Psychophysics, 63(8), 1293–1313. https://doi.org/10.3758/BF03194544

Winter, D. A. (2009). Biomechanics and motor control of human movement (4th ed). Wiley.

